# Normal observers show no evidence for blindsight in facial emotion perception

**DOI:** 10.1101/314906

**Authors:** Sivananda Rajananda, Jeanette Zhu, Megan A.K. Peters

## Abstract

It is commonly assumed that normal human observers can exhibit ‘blindsight-like’ behavior: the ability to discriminate or identify a stimulus without being aware of it. However, we recently used a bias-free task to show that what looks like blindsight may in fact be an artifact of typical experimental paradigms’ susceptibility to response bias. While those findings challenge many previous reports of blindsight in normal observers, they do not rule out the possibility that different stimuli or techniques could reveal such perception without awareness. One particularly intriguing candidate for this unconscious perception is emotion processing, as processing of emotional stimuli (e.g. fearful/happy faces) has been reported to potentially bypass conscious visual circuits. Here we used the bias-free blindsight paradigm to investigate whether emotion processing might reveal ‘featural blindsight’, i.e. the ability to identify a face’s emotion without having introspective access to the task-relevant features of the face that led to the discrimination decision. However, despite the purported ability of emotional stimuli to bypass conscious visual processing, we saw no evidence for such emotion processing ‘featural blindsight’: like in our previous study, as soon as participants could identify a face’s emotion they reported introspective access to the task-relevant features, matching predictions of a Bayesian ideal observer. The present results challenge dominant theory, adding to the growing body of evidence that perceptual discrimination ability in the complete absence of introspective access may not be possible for neurologically intact observers.

---

The neurological condition of blindsight has fascinated consciousness researchers since its discovery (Weiskrantz, 1986, 1996); in this rare condition, patients with damage to primary visual cortex can directly discriminate some aspects of visual stimuli but report no conscious visual awareness of them. Thus, to experimentally use this dissociation to isolate conscious awareness of a stimulus from the potential confound of signal processing capacity (Giles, Lau, & Odegaard, 2016; Lau, 2008; Peters, Kentridge, Phillips, & Block, 2017), we must find a way to induce blindsight-like behavior in neurologically intact observers. Successful induction of such “performance without awareness” would pave the way for computational and neuroimaging studies seeking to identify the neural correlates of consciousness (Aru, Bachmann, Singer, & Melloni, 2012; Block, 2005; Rees, Kreiman, & Koch, 2002; Tononi & Koch, 2008).

Unfortunately, most attempts to induce blindsight-like behavior in normal observers -- typically using visual masking or other similar techniques -- are susceptible to response bias confounds (Charles, King, & Dehaene, 2014; Charles, Van Opstal, Marti, & Dehaene, 2013; Eriksen, 1960; Hannula, Simons, & Cohen, 2005; Jachs et al., 2015; Lloyd, Abrahamyan, & Harris, 2013; Merikle, Smilek, & Eastwood, 2001; Ramsøy & Overgaard, 2004). Further, past attempts to remedy response bias concerns (Kolb & Braun, 1995; Kunimoto, Miller, & Pashler, 2001) have met with their own conceptual or replicability challenges (Evans & Azzopardi, 2007; Galvin, Podd, Drga, & Whitmore, 2003; Morgan, Mason, & Solomon, 1997; Robichaud & Stelmach, 2003). Recently, we used visual masking in a bias-free paradigm to demonstrate that it may not be possible to induce blindsight in normal observers when response bias confounds are controlled for (Peters & Lau, 2015); follow-up studies also demonstrated that several other masking techniques commonly assumed to dissociate objective and subjective processing may similarly fail to produce conditions under which blindsight could be induced (Knotts, Lau, & Peters, 2018).

Yet these failures do not unequivocally prove that blindsight-like behavior *cannot* be induced in normal observers. Although we failed to demonstrate that normal observers can discriminate a stimulus yet be completely unaware of its *overall presence* (Peters & Lau, 2015), it may be possible for an observer to be able to discriminate a stimulus above chance while being unaware of its *task-relevant properties*. This might occur especially for stimuli that could possibly bypass ‘conscious’ visual processing areas to activate other task-relevant circuitry, such as observations of amygdala reactivity in ‘unconscious’ face emotion processing (Diano, Celeghin, Bagnis, & Tamietto, 2016; Khalid & Ansorge, 2017; Pessoa & Adolphs, 2010; Watanabe & Haruno, 2015). In such emotion processing ‘featural blindsight’, the observer would be aware of the face itself and be able to correctly identify its emotion (e.g. happy versus fearful), but would report no subjective or introspective access to the *task-relevant features*, evinced by no confidence in their choices. Indeed, it has recently been reported that processing of emotional stimuli, especially fearful stimuli, may be a powerful candidate for inducing blindsight-like behavior (Vieira, Wen, Oliver, & Mitchell, 2017).

Therefore, here we used emotional face stimuli matched on all low-level properties (e.g. luminance, contrast, spatial frequency) to investigate whether normal observers can identify facial emotions without being aware of their ability to do so. Importantly, we controlled for response bias using the same bias-free paradigm previously shown to demonstrate optimal introspective access using low-level stimuli (i.e. masked Gabor patches) (Peters & Lau, 2015). If we observe emotion-processing ‘featural blindsight’ using this paradigm, the results would suggest that the previous failure to demonstrate blindsight in normal observers was due to the unsuitability of impoverished, low-level stimuli to dissociate objective versus subjective thresholds (Peters & Lau, 2015) -- and further, that emotion processing may provide an ideal candidate for experimentally isolating subjective awareness from objective task performance capacity in future studies seeking the neural or computational correlates of consciousness.

## Methods

### Behavioral Experiment

#### Participants

Participants for this experiment were recruited from Amazon Mechanical Turk. The task was launched for 40 people; 33 of these participants completed the experiment. Those who completed the experiment were paid $2. Participants were also incentivized to perform well with an extra $1 which was awarded if they performed better than the previous participant (incentive structure is described under *Procedure*). Of the 33 people who completed the experiment, 2 had incomplete data, and 2 had >20% of trials which met the exclusion criteria (see below); data from the remaining 29 participants was included in all further analyses. Informed consent was obtained before the start of the experiment, and all procedures were approved by the University of California, Riverside Institutional Review Board and made in accordance with the Declaration of Helsinki.

#### Stimuli

Stimuli consisted of 4 sets of male and 4 sets of female faces displaying happiness (‘happy’ faces), fear (‘fearful’ faces), or no emotion (‘neutral’ faces). The faces were selected from the NimStim set of facial expressions (Tottenham et al., 2009), which contains images of 43 (18 male, 25 female) people displaying various emotions and neutral expressions. In an earlier experiment (also via Amazon Mechanical Turk), 20 participants viewed 15 female faces and 13 male faces displaying happiness and fear at maximal intensity, and rated the intensity of the emotion shown by each face on a scale of 0-100, where 0 is “completely neutral” and 100 is the “most possible fear/happiness that a person could exhibit”. We selected the 4 highest-rated faces from each gender for which the difference in the emotional intensity ratings was not significantly different for ‘happy’ versus ‘fearful’ images.

The selected 4 male and 4 female faces were then morphed between the ‘neutral’ image and the ‘happy’ and ‘fearful’ images for each person, to create a set of 8 morphs from 0% emotional intensity to 100% emotional intensity for each emotion. We then matched the low-level image properties of the morphs using the SHINE toolbox (Willenbockel et al., 2010): image luminance histograms of the photos were matched through SHINE toolbox’s histMatch function with the structural similarity also optimized through SSIM gradient ascent (20 iterations), and the amplitude spectrum were matched across images through the specMatch function.

#### Procedure

On each trial, subjects viewed two faces presented sequentially, one after the other. Following the 2-interval forced choice (2IFC) confidence-betting paradigm used by Peters and Lau (2015), one interval showed the ‘neutral’ face (Emotion-Absent interval; EA) and the other showed the same person’s face but with some degree of emotion, selected from the morphed images (Emotion-Present interval; EP). Subjects were asked to determine the emotion of each face (happy/fearful); they were not told that one face by definition showed neutral emotion. Subjects were also instructed that the two intervals’ emotions were independently chosen, such that any combination of emotion pairs was possible: happy/happy, happy/fearful, fearful/happy, fearful/fearful. In addition to identifying the emotion, subjects were asked to bet on which emotional identification they felt more confident in. To incentivize their performance, they were told that for each emotion that they identified correctly, they would get a point. They were also told that if they bet on the easier interval, i.e. the one they thought they were more likely to get correct, they would also get a point. These points were then used to determine if they earned the bonus payment.

Each trial began with a fixation point (1000ms) followed by the first interval’s face presented centrally (size dynamically scaled based on window size to occupy approximately 60% of screen height) (500ms), a second fixation point (1000ms), and the second interval’s face presented centrally (500ms) (Figure 1). The order of EP and EA intervals was counterbalanced across trials, and the emotion to be presented in the EP interval was chosen pseudorandomly such that 50% of trials presented a fearful face and 50% of trials presented a happy face in the EP interval. Each of the two faces used on each trial was chosen randomly from the 8 possible face sets, and both faces within a trial were from different sets. For the EP interval in each trial, the emotional intensity of the face (whether happy or fearful) was pseudorandomly set to be 5%, 15%, or 25% for 32 trials each, or 75% for 16 trials (75% is very easy).

**Figure 1.**
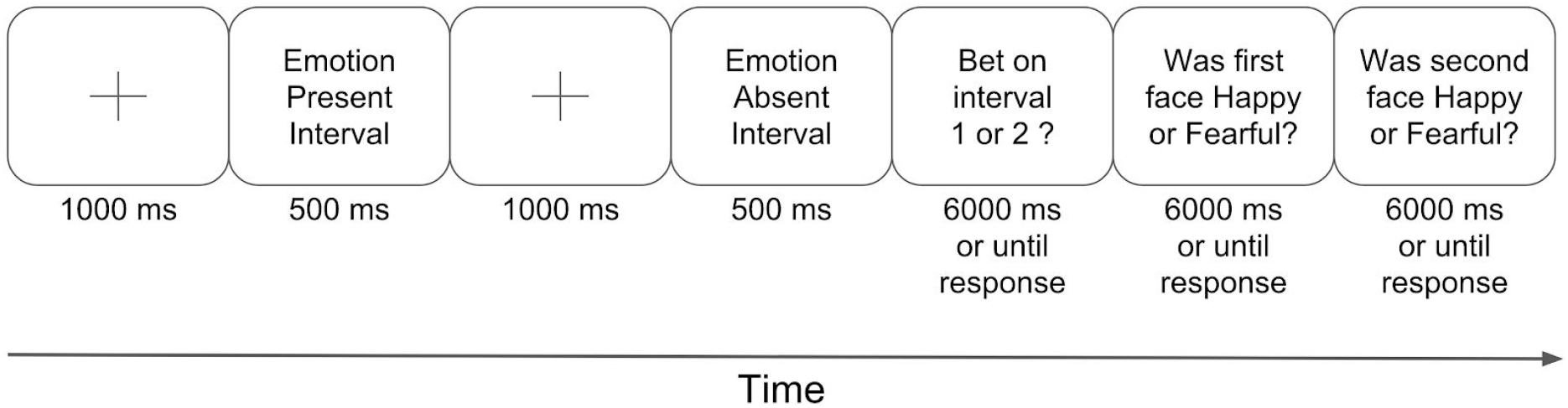
A sample 2-interval forced-choice (2IFC) trial. Subjects viewed two intervals containing a face with Emotion Present (EP interval) or Emotion Absent (EA interval), then ‘bet’ on which emotion discrimination they thought they were more likely to get correct. Subjects then indicated the emotion they identified in the first and second intervals, respectively. The order of the Emotion Present (EP) and Emotion Absent (EA) intervals was counterbalanced between trials.

After the second interval’s face had disappeared, subjects indicated which interval they felt more confident in (Interval Question), and then the emotion decision (Emotion Questions) for the first and second face, respectively. To indicate their answers, subjects entered their ‘emotion’ choices using the “H” key (happy) and “F” key (fearful) on their keyboard using their right hand, and their choices about which decision to ‘bet’ on using the “1” and “2” keys using their left hand. The next trial began as soon as subjects had pressed a key or after 6s had elapsed, whichever happened sooner. As described above, only one of the intervals contained an emotional face, with the other containing a neutral face -- despite the fact that subjects were told both intervals contained emotional faces.

Before beginning the main experiment, participants completed 4 easy practice trials and 4 hard practice trials. Practice trials were identical to trials in the main experiment, except that the faces presented in both intervals were presented for 1000ms instead of 500ms, and *both* faces had an emotion present to reinforce the instructions given to subjects to judge the emotion in *both* intervals.The emotion intensity in the easy practice trials was 100%, while the emotion intensity in the hard practice trials was 25%. Prior to the beginning of the hard practice trials, participants were told that some emotions would be hard to distinguish, and if they were unsure, they should just make their best guess. After the 8 practice trials, the participants completed 112 trials in the main experiment.

Participants were given two breaks: one after trial 37, and another after trial 74. Each break lasted 1 minute, although participants were encouraged to take more time if they needed. In total, the experiment lasted approximately 30 minutes.

#### Analysis

To determine the objective (Type 1) performance capacity for each participant, we calculated the percent of EP trials in which the emotion was correctly identified separately for each emotional intensity level (Emotion Questions). This produced four ‘% correct emotion discrimination’ values for each subject. Following Peters & Lau (2015), we also calculated the confidence/introspective ability of each observer (Type 2) as the percent of trials the subject ‘bet’ on the EP interval at each of the four emotional intensity levels (Interval Question; ‘% bet on EP’). Trials were excluded from the ‘% bet on EP’ analysis if reaction times to any question (Emotion or Interval Questions) in the trial exceeded 4000ms, or if the subject failed to respond to all questions in the trial. Likewise, trials were excluded from the ‘% correct’ analysis if reaction time to the Interval Question or the Emotion Question for the EP interval exceeded 4000ms, or if the subject failed to respond to either of these questions in the trial. We excluded from further analysis any subjects who retained fewer than 80% valid trials after the two exclusion criteria above were implemented.

To look for emotion processing ‘featural blindsight’, we follow the definitions introduced by Peters and Lau (2015). If subjects show no emotion processing ‘featural blindsight’, as soon as they are able to meaningfully identify the emotion in the EP interval (‘% correct emotion discrimination’ > 50%), they should exhibit some degree of introspective access evinced by an ability to bet on their choices (‘% bet on EP interval’ > 50%); this behavior would match predictions of a Bayesian ideal observer (Peters & Lau, 2015). If, in contrast, subjects do exhibit ‘featural blindsight’, there will be some level of objective performance capacity (‘% correct emotion discrimination’ > 50% in the EP interval) at which subjects have *no* introspective access to their decisional processes, i.e. *no* ability to meaningfully bet on their choices (‘% bet on EP interval’ = 50%). To arbitrate between these hypotheses, we compare the goodness of fit of a Bayesian ideal observer computational model to that of a Bayesian observer with additional Type 2 noise, which exhibits featural blindsight. See next sections for details.

During the calculation of d’, whenever the subject exhibited a hit rate or false alarm rate of 0 or 1, we applied a standard correction (Wickens, 2001) to ensure that the calculated d’ would be within a reasonable range (i.e. not infinity). For the Hit Rate (HR), we corrected the probability using the formulas

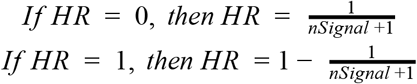

where *nSignal* is the number of emotion present intervals for that emotional intensity. For the False Alarm Rate (FAR), we corrected the probability using the formulas

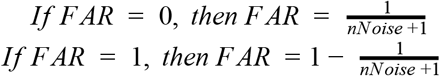
where *nNoise* is the number of emotion present intervals for that emotional intensity.

### Computational Model

The Bayesian ideal observer computational model has previously been described elsewhere (Peters & Lau, 2015). Briefly, for every 2IFC trial, we draw two samples *d_EP_* and *d_EA_*, each from a bivariate Gaussian distributions *S_EP_* and *S_EA_* (*S ~ N*(*μ*, *Σ*)) representing an Emotion-Present face and an Emotion-Absent face, respectively. The mean of the generating distribution for the EP face is *μ_EP_* = [*e_Happy_*, 0] or *μ_EP_* = [0, *e_Fearful_*], depending on the random assignment of emotional valence in the EP interval; the mean of the generating distribution for the EA face is always *μ_EA_* = [0, 0]. *Σ* = [1 0; 0 1], the identity matrix, for all generating distributions.

In all cases, the true emotional intensity that generated the emotion, i.e. *e_emotion_*, is unknown to the observer; the observer only has access to the sample that is being observed. This is why the decision about whether each sample is happy or fearful is made according to Bayes’ rule, marginalizing across all possible emotional intensity levels for both Happy and Fearful emotions:

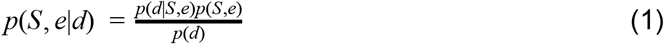

Confidence in the decision (happy or fearful) for each interval is defined as the posterior probability of the choice that was made, i.e.

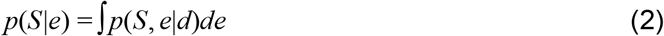

The chosen emotion for each interval, both EP and EA, is defined as the emotion *i* that maximizes this posterior probability, i.e.

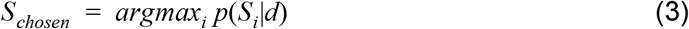

The model then ‘bets’ on the interval that has the higher posterior probability of the choice, i.e. higher confidence, by comparing the posterior probabilities for the chosen emotion in the EP and EA intervals via a decision variable *D*,

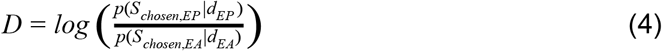

If *D* ≥ 0, the model ‘bets’ on the EP interval; if D < 0, the model ‘bets’ on the EA interval.

All simulations of this model’s behavior were done via custom-written scripts in Matlab (Natuck, MA), with emotional intensity level *e* ranging from 0.01 to 1 in steps of 0.01 and 5000 samples per emotional intensity level.

#### Simulating emotion processing ‘featural blindsight’

The ideal observer described above has access to Type 2 information (confidence) with the same degree of fidelity as its access to Type 1 information (emotion decision). In contrast, an observer that exhibits blindsight would have much poorer access to Type 2 information than Type 1 information. Following previous work (Maniscalco & Lau, 2016; Peters & Lau, 2015), this can be modeled as the addition of increasing amounts of Type 2 noise after the Type 1 decision has been reached. To simulate the behavior of such an observer, we therefore added Gaussian noise to the definition of the Type 2 decision variable *D* after the Type 1 decision, i.e.

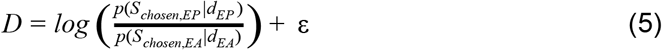

where *ε* ~ *N*(0, *σ*). We simulated predicted behavior for increasing degrees of ‘featural blindsight’ at values of *σ* from 0.01 to 1 in steps of 0.01.

#### Goodness of fit

The goodness of fit of both models -- the Bayesian ideal observer and the ‘featural blindsight’ observer at various levels of Type 2 noise (Eq. 5) -- was calculated as the multinomial log-likelihood (*L_m_*) of the model with parameters *Φ* given the data. This measure quantifies the relative agreement between the data collected from participants and that predicted by a model. We use the following formula:

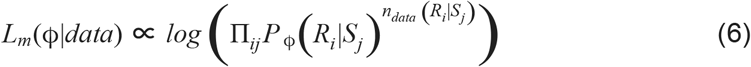

where *S_j_* is the type of stimulus that might be shown on a given trial, and *R_i_* refers to the behavioral response a subject produces on that trial. *n_data_*(*R_i_|S_j_*) represents the count of how many times a human observer produced response *Ri* after viewing stimulus *S_j_*. P_Φ_(*R_i_|S_j_*) represents the model’s prediction of the probability a subject produced response *Ri* after viewing stimulus *Sj*, according to the model with parameters *Φ*, i.e. the percentage of time the model produced this ‘response’ to this ‘stimulus’. Importantly, the multinomial log-likelihood quantifies the fit of the *full* distribution of probabilities of each response type given each stimulus type, not just with reference to a summary statistic.

We used this metric to evaluate the predictions of both models at all levels of Type 1 performance exhibited by each human observer. Note that for the ideal observer, there are no free parameters; for the ‘featural blindsight’ observer with Type 2 noise, the only parameter *ϕ* is the magnitude of the Type 2 noise (Eq. 5).

## Results

Two subjects had incomplete data and our exclusion criteria led us to exclude another two subjects from our analyses, leaving 29 subjects. Across subjects, the four levels of emotion presented typically led to above-chance performance for each emotion intensity: 5% (mean % correct = 49.92 ± SD 0.0535), 15% (mean % correct = 0.6132 ± SD 0.0908), 25% (mean % correct = 0.7009 ± SD 0.1056), and 75% (mean % correct = 0.8971 ± SD 0.1385). A one-way repeated measures ANOVA showed the expected increase in Type 1 performance (‘% correct emotion discrimination’) as a function of increasing emotional intensity (F(2.386) = 105.223, p <.001, Greenhouse-Geisser corrected for sphericity violation). Likewise, subjects’ ability to bet on the EP interval also increased with increasing emotional intensity as expected (F(2.112) = 35.840, p <.001, Greenhouse-Geisser corrected for sphericity violation).

The critical analysis is whether subjects show emotion processing ‘featural blindsight’. Following the definitions introduced by Peters and Lau (2015), if subjects show such emotion processing ability without awareness, we would expect that there would be some level of objective performance capacity (‘% correct emotion discrimination’ > 50% in the EP interval) at which subjects had no ability to meaningfully bet on their choices (‘% bet on EP interval’ = 50%). To look for this possibility, we plotted the ‘% bet on EP interval’ values against the ‘% correct emotion discrimination’ values for each subject, and fit a smoothing spline (using Matlab’s smoothingspline function, which minimizes the expression 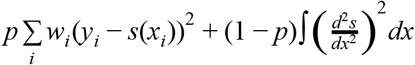, with the smoothing parameter set to 0.99) to the data across all subjects (Figure 2a). Data from individual subjects closely resembles the group data (Supplementary Figure S1).

**Figure 2.**
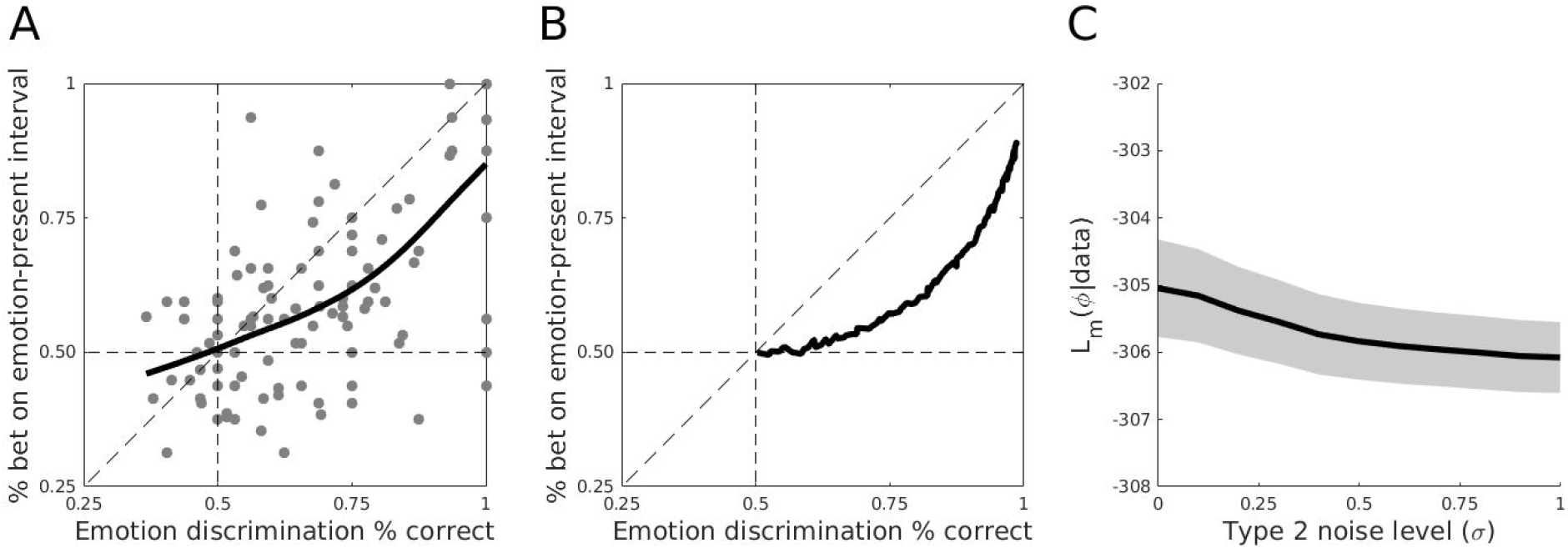
Human subjects show no emotion processing ‘featural blindsight’. (a) As soon as subjects are able to correctly identify the emotion of the face in the Emotion Present (EP) interval (‘% correct emotion discrimination’ > 50%), they appear to be able to meaningfully bet on their choices (‘% bet on EP interval’ > 50%). (b) Subjects’ data closely matches predictions from the Bayesian ideal observer computational model. (c) Increasing levels of Type 2 noise (*σ*; see Methods) to produce emotion processing ‘featural blindsight’ in the computational model leads to increasingly *worse* goodness of fit (*L_m_*) between the model and the human subjects’ data (F(1.141) = 15.541, p <.001, Greenhouse-Geisser corrected for sphericity violation; two-tailed paired samples t-test between extremes of *σ* = 0 [no Type 2 noise] and *σ* = 1 [large Type 2 noise]: t(28) = 3.9255, p = < .001). Error cloud represents the standard error of the mean.

From visual inspection alone, we see no evidence of emotion processing ‘featural blindsight’: as soon as subjects are able to correctly identify the emotion in the EP interval, they are able to meaningfully bet on their choices to some degree. The pattern of the data closely mimics the failure to observe blindsight-like behavior in normal observers using low-level visual stimuli, in which the target-absent interval (here the EA interval) contained no stimulus at all (Peters & Lau, 2015). It also visual matches predicted behavior from a Bayesian ideal observer (Figure 2b).

However, because this study was done online through Amazon mTurk, the data for individual subjects is somewhat noisier than it would be had subjects each completed many hours of psychophysics, as they did in the original Peters and Lau (2015) study (Supplementary Figure 1). Therefore, we rely on the quantitative goodness of fit metrics to critically evaluate whether any hint of ‘featural blindsight’ might be present.

We calculated the goodness of fit as *L_m_* for each human observer for the Bayesian ideal observer, and compared it to *L_m_* for the ‘featural blindsight’ observer model at each level of Type 2 noise (see Methods, Eq. 6). This analysis revealed that as Type 2 noise increases -- i.e. as ‘featural blindsight’ becomes stronger -- the model fits the data less and less well (main effect of Type 2 noise magnitude, F(1.141) = 15.541, p <.001, Greenhouse-Geisser corrected for sphericity violation; Figure 2c). This interpretation was confirmed with a two-tailed paired samples t-test of the log likelihoods between *σ* = 0 (no Type 2 noise, i.e. Bayesian ideal observer) and *σ* = 1 (high Type 2 noise) confirms that goodness of fit decreases as Type 2 noise increases (t(28) = 3.9255, p = < .001). Thus, the Bayesian ideal observer with no Type 2 noise -- i.e. the observer with no ‘featural blindsight’ -- provides the best explanation of the human subjects’ behavior.

## Discussion

Here we used a bias-free two-interval forced-choice (2IFC) method to examine whether facial emotion processing might reveal blindsight-like behavior in normal human observers. We modified a previous version of this paradigm (Peters & Lau, 2015) to match low-level visual properties such as luminance, contrast, and spatial frequency differences between the Emotion Present (EP) and Emotion Absent (EA) intervals; our goal was to reveal emotion processing ‘featural blindsight’, in which observers would be able to correctly identify an emotion when it was present but have no ability to introspect on the evidence or process leading to their choices. However, despite previous reports suggesting that emotion processing may bypass conscious visual processing areas (Diano et al., 2016; (Khalid & Ansorge, 2017; (Pessoa & Adolphs, 2010; (Watanabe & Haruno, 2015) and that emotional (especially fearful) stimuli may be powerful candidates for inducing blindsight-like behavior (Vieira et al., 2017), here we saw no evidence to suggest that emotion-processing ‘featural blindsight’ may occur.

These results are in line with the absence of evidence for blindsight-like behavior reported in the original Peters and Lau (2015) study (which used forward-backward masking and simple visual stimuli), as well as the observation that various masking techniques fail to produce differences in the relationship between objective versus subjective thresholds, contrary to dominant theory (Knotts et al., 2018). Our observations also support the finding that even noninvasive transcranial magnetic stimulation fails to produce the ability to discriminate a stimulus in the absence of any visual awareness or confidence (Peters, Fesi, et al., 2017). Taken together, the evidence appears to be mounting that it is extremely difficult -- if not impossible -- to use visual masking or noninvasive techniques to produce dissociations between performance and awareness that would be ideal for experimentally isolating subjective consciousness.

However, despite the consistency of the present results with previous reports, it should certainly be noted that the data collection method we used here was quite different from the standard approach. Each of our subjects completed only 112 trials in the main experiment across four levels of emotion intensity, which is significantly less than the volume of data collected in e.g. the original study (2600 trials; (Peters & Lau, 2015)). One could argue that the online data collection method, with so few trials per subject, may preclude the possibility of detecting the blindsight-like effect we hoped to reveal. In response, we note that each of our 33 subjects’ data looks much like the data in aggregate, and that the goodness of fit metrics were calculated for each subject individually rather than for the group average. That we identified decreasing goodness of fit for increasing Type 2 noise, despite the noisiness of online data collection and the small trial numbers, suggests that the data collection method was not so noisy as to prevent the possibility of identifying emotion processing ‘featural blindsight’. Future studies may wish to follow up the results presented here with additional variations, done both in the laboratory and in an online setting.

Another way to check whether our online data collection method produces meaningful data would be to compare the emotional discrimination results to previous reports. In another study of emotion processing and confidence, it was reported that fear processing is ‘special’: when perceiving a fearful face human subjects show a liberal bias in both detection and discrimination tasks, and ‘fear’ choices are also reported with higher confidence (Koizumi, Mobbs, & Lau, 2016). We used signal detection theoretic approaches (Green & Swets, 1966; Macmillan & Creelman, 2004) to examine whether the same biases would be present in our own data. Consistent with previous reports that fearful faces are perceived more easily or quickly (Amting, Greening, & Mitchell, 2010; Milders, Sahraie, Logan, & Donnellon, 2006; Phelps, Ling, & Carrasco, 2006; Stein, Peelen, Funk, & Seidl, 2010; Stein, Seymour, Hebart, & Sterzer, 2014; Stein, Zwickel, Ritter, Kitzmantel, & Schneider, 2009; Stienen & de Gelder, 2011), we also saw a significant shift in the Type 1 decisional criterion that biased subjects to respond ‘fearful’ more than ‘happy’ at lower emotional intensities in both the EP (t_5%_(28) = 4.174, p < .001; t_15%_(28) = 3.917, p < .001; t_15%_(28) = 2.040, p = .051; 75% intensity N.S.; 5% and 15% intensities are significant with Bonferroni correction for multiple comparisons) and EA (t(28) = 5.2189, p < .001) intervals. Interestingly, the magnitude of this shift when significant or trending (*c_EA_* = −0.600, *c_5%_* = −0.444, *c_15%_* = −0.352, *c_15%_* = −0.187) was somewhat larger than that reported by Koizumi and colleagues (2016) in their discrimination task (*c* ≈ −0.12). This result provides further evidence that online data collection, despite being impoverished relative to the controlled setting of the laboratory, may be a viable method for identifying even small effects.

The present results -- and previous studies showing similar failure to induce blindsight-like behavior using generally accepted methods for doing so (Peters, Fesi, et al., 2017; Peters & Lau, 2015) -- present a growing challenge to the dominant view that stimulus manipulation and/or noninvasive brain stimulation may result in task performance capacity in the absence of introspective access (Boyer, Harrison, & Ro, 2005; Charles et al., 2013; Kolb & Braun, 1995; Merikle, 1982; Merikle et al., 2001; Reingold & Merikle, 1988). We acknowledge that the growing body of work we add to here does not speak definitively to the *impossibility* of inducing dissociations between the objective and subjective thresholds in normal human observers. However, these results do increasingly suggest that inducing blindsight-like perceptual capacity in the absence of introspective access may be much more difficult to induce than commonly believed. Consequently, demonstrating such a dissociation using this conservative 2IFC confidence method would provide indisputable evidence that blindsight *can* be induced in normal observers if one employs the appropriate experimental manipulations.

## Data accessibility statement

The code used in the simulations and data presented here can be found at https://github.com/vrsivananda/Faces_2IFC_Task.

## Context paragraph

How can we experimentally isolate ‘consciousness’, in order to study it scientifically? One of the most promising avenues has been to induce ‘blindsight’-like behavior in normal human observers: by manipulating the stimulus or experimental design, it might be possible to make people unconscious of a stimulus but still be able to identify it. However, recent studies have suggested that maybe such ‘unconscious perception’ is just response bias, i.e. people saying they cannot see a stimulus when it seems very dim or uncertain. Here we explored whether emotion perception might be an ideal candidate for inducing blindsight-like behavior, because it is thought to rely on circuits that can bypass conscious visual areas. However, as in previous experiments using lower level stimuli, we found no evidence for perception without awareness. This finding adds to a growing pool of evidence from bias-free experimental paradigms suggesting that perception without awareness might not exist, or at least might be much harder to induce than previously thought. We hope that via increased adoption of these bias-free paradigms, we will be able to promote the study of consciousness in a rigorous manner.

## Supplemental Material

Normal human observers show no blindsight in facial emotion perception Sivananda Rajananda, Jeanette Zhu, Hakwan Lau, & Megan A. K. Peters

---

**Figure S1.**
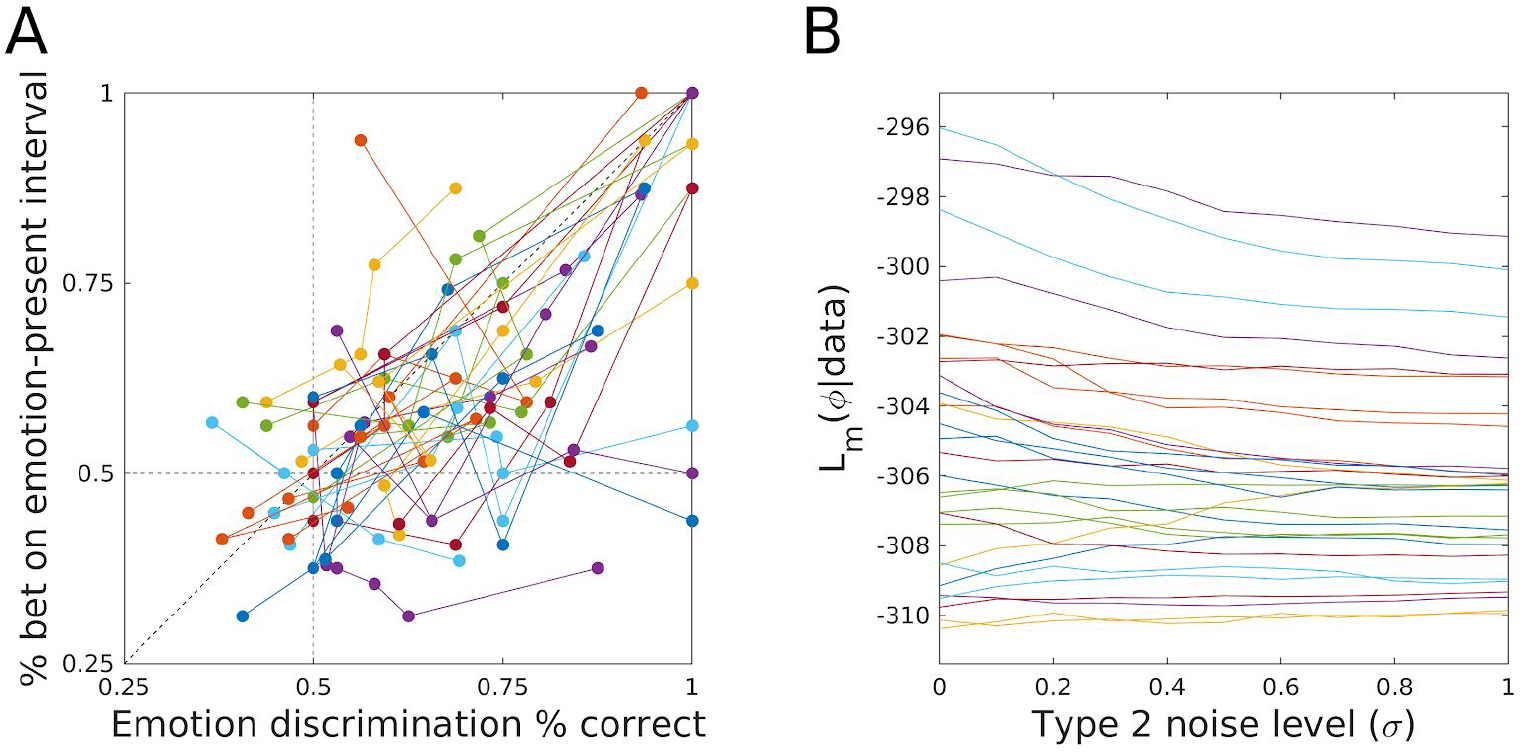
Individual subjects’ behavioral results and goodness of fit. (a) Individual subjects’ results closely mimic aggregate results, demonstrating no convincing evidence for emotion processing ‘featural blindsight’. As soon as subjects are able to discriminate the Emotion Present interval’s emotion above chance, they have some ability to bet on their choices, as predicted by a Bayesian ideal observer. (b) As Type 2 noise increases for the ‘featural blindsight’ observer model, the goodness of fit of the model to the data decreases. This shows that of the models we tested, the Bayesian ideal observer with no ‘featural blindsight’ provides the best fit to each individual subject’s behavior. In both panels, each color represents one individual subject’s data.

**Figure S2.**
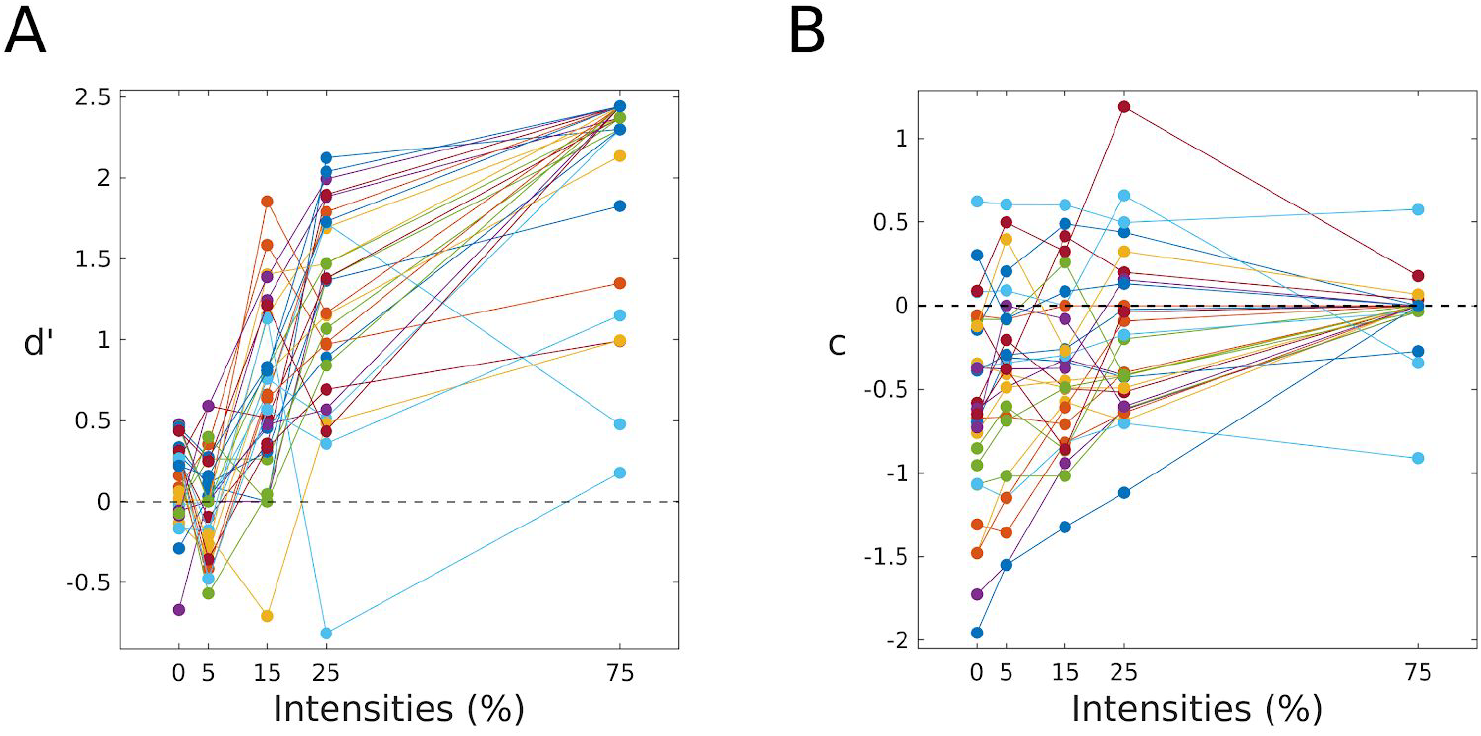
Type 1 performance (d’) and response bias (*c*) as a function of stimulus emotion intensity. (a) As emotion intensity increases, subjects show the expected increase in objective performance capacity. (b) Subjects exhibit significant bias to respond ‘fearful’, especially at low emotion intensities. As above, in both panels, each color represents one individual subject’s data.

